# Immunogenicity of the BA.5 Bivalent mRNA Vaccine Boosters

**DOI:** 10.1101/2022.10.24.513619

**Authors:** Ai-ris Y. Collier, Jessica Miller, Nicole P. Hachmann, Katherine McMahan, Jinyan Liu, Esther Apraku Bondzie, Lydia Gallup, Marjorie Rowe, Eleanor Schonberg, Siline Thai, Julia Barrett, Erica N. Borducchi, Emily Bouffard, Catherine Jacob-Dolan, Camille R. Mazurek, Audrey Mutoni, Olivia Powers, Michaela Sciacca, Nehalee Surve, Haley VanWyk, Cindy Wu, Dan H. Barouch

## Abstract

Waning immunity following mRNA vaccination and the emergence of SARS-CoV-2 variants has led to reduced mRNA vaccine efficacy against both symptomatic infection and severe disease. Bivalent mRNA boosters expressing the Omicron BA.5 and ancestral WA1/2020 Spike proteins have been developed and approved, because BA.5 is currently the dominant SARS-CoV-2 variant and substantially evades neutralizing antibodies (NAbs). Our data show that BA.5 NAb titers were comparable following monovalent and bivalent mRNA boosters.

---

Waning immunity following mRNA vaccination and the emergence of SARS-CoV-2 variants has led to reduced mRNA vaccine efficacy against both symptomatic infection and severe disease^1,2^. Bivalent mRNA boosters expressing the Omicron BA.5 and ancestral WA1/2020 Spike proteins have been developed and approved, because BA.5 is currently the dominant SARS-CoV-2 variant and substantially evades neutralizing antibodies (NAbs)^3^. However, the immunogenicity of the BA.5-containing bivalent mRNA boosters remains unknown.

We evaluated humoral and cellular immune responses in 15 individuals who received the original monovalent mRNA boosters and in 18 individuals who received the bivalent mRNA boosters (**Table S1**). Participants had a median of 3 (range 2-4) prior COVID-19 vaccine doses, and 33% had documented SARS-CoV-2 infection during the Omicron surge, although it is likely that the majority of participants had hybrid immunity prior to boosting given the high prevalence and limited severity of Omicron infection. Both the monovalent and bivalent mRNA boosters led to preferential expansion of WA1/2020 NAb titers and lower BA.1, BA.2, and BA.5 NAb titers (**Fig. 1A, B**). Median BA.5 NAb titers increased from 184 to 2,829 following monovalent mRNA boosting and from 211 to 3,693 following bivalent mRNA boosting. The Pfizer and Moderna bivalent mRNA boosters induced similar NAb profiles (**Fig. S1**). Binding antibody responses by ELISA and electrochemiluminescence assays were comparable following monovalent and bivalent mRNA boosting (**Fig. S2, S3**).

**Figure 1.**
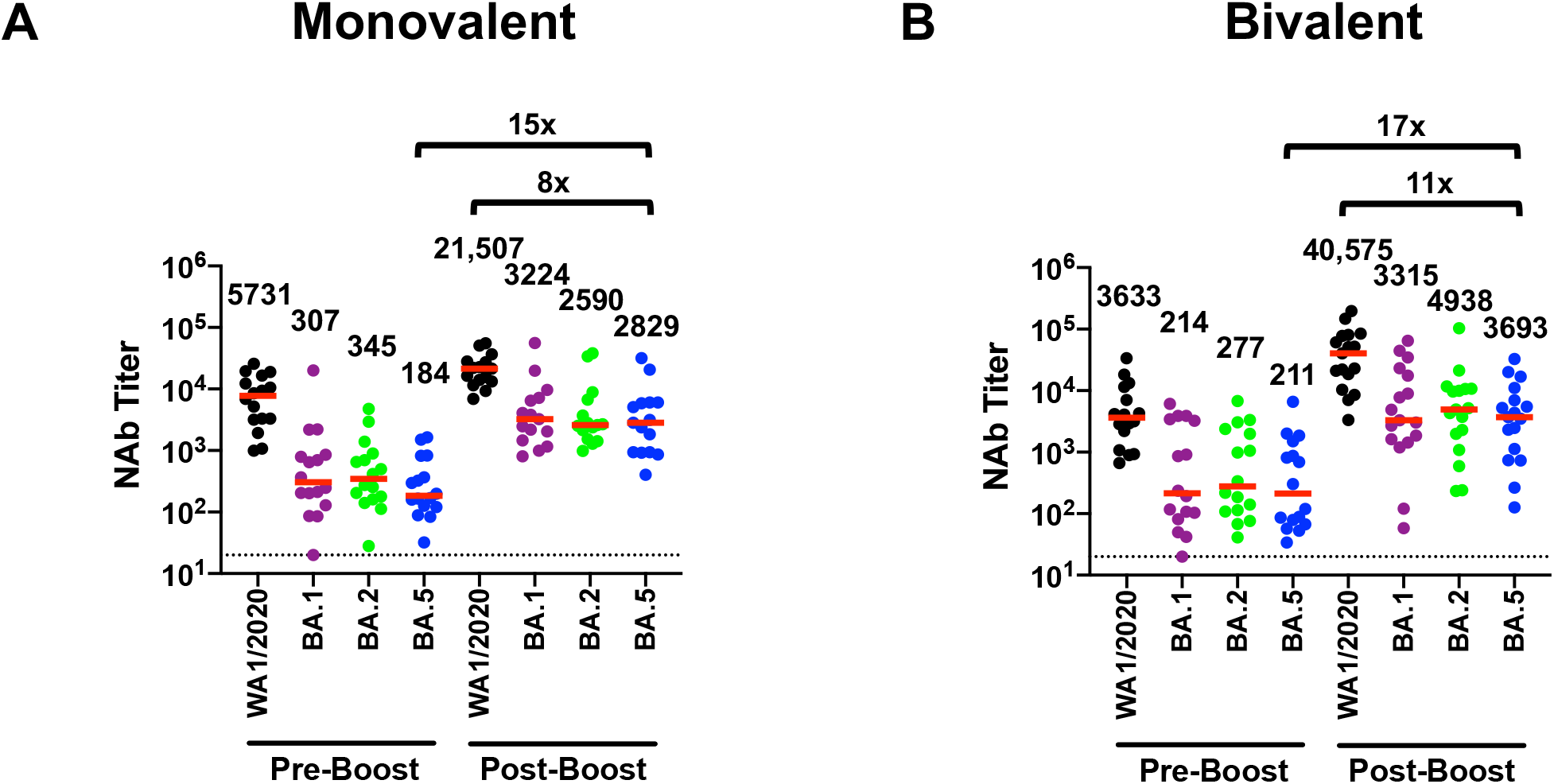
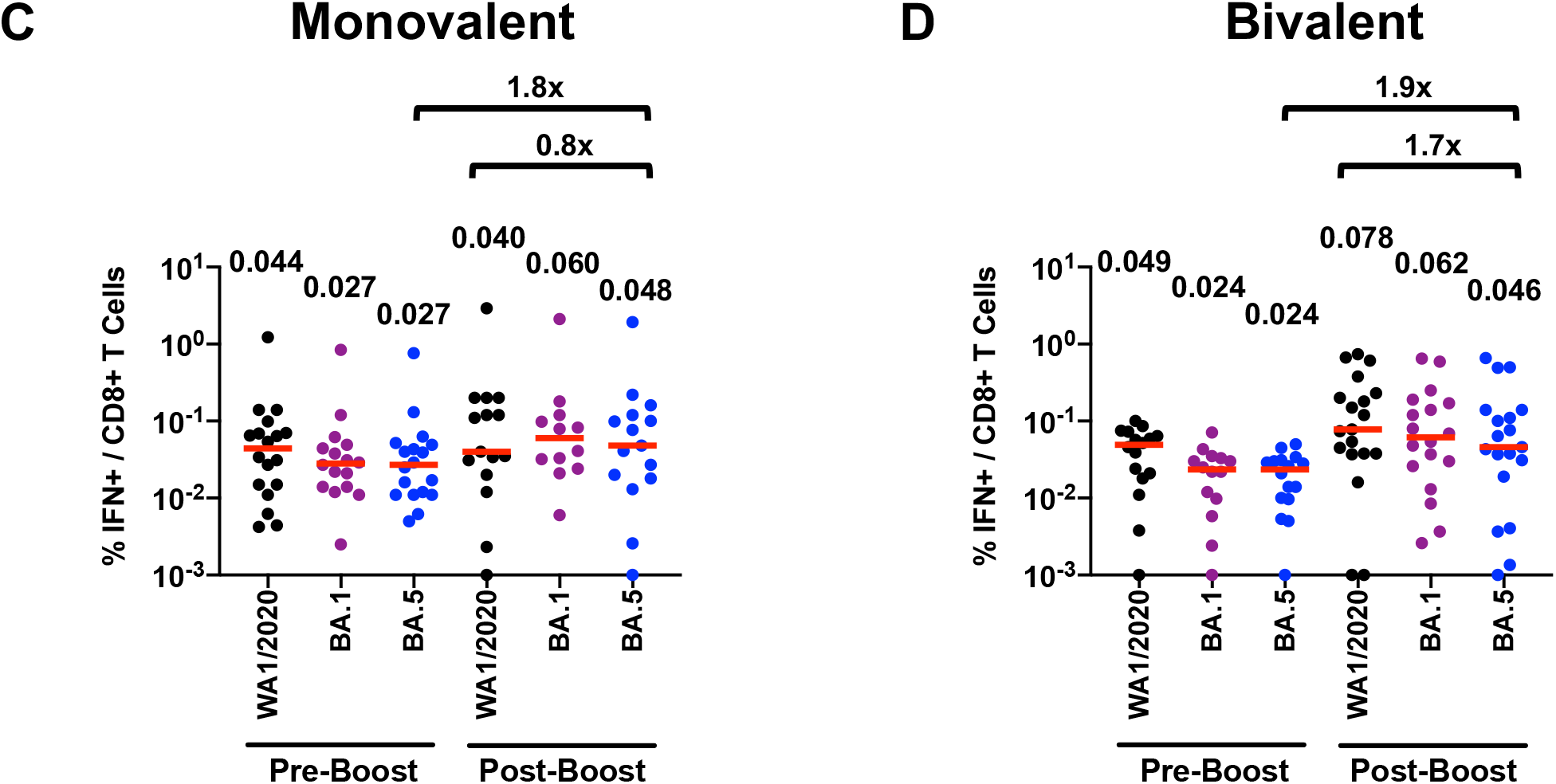
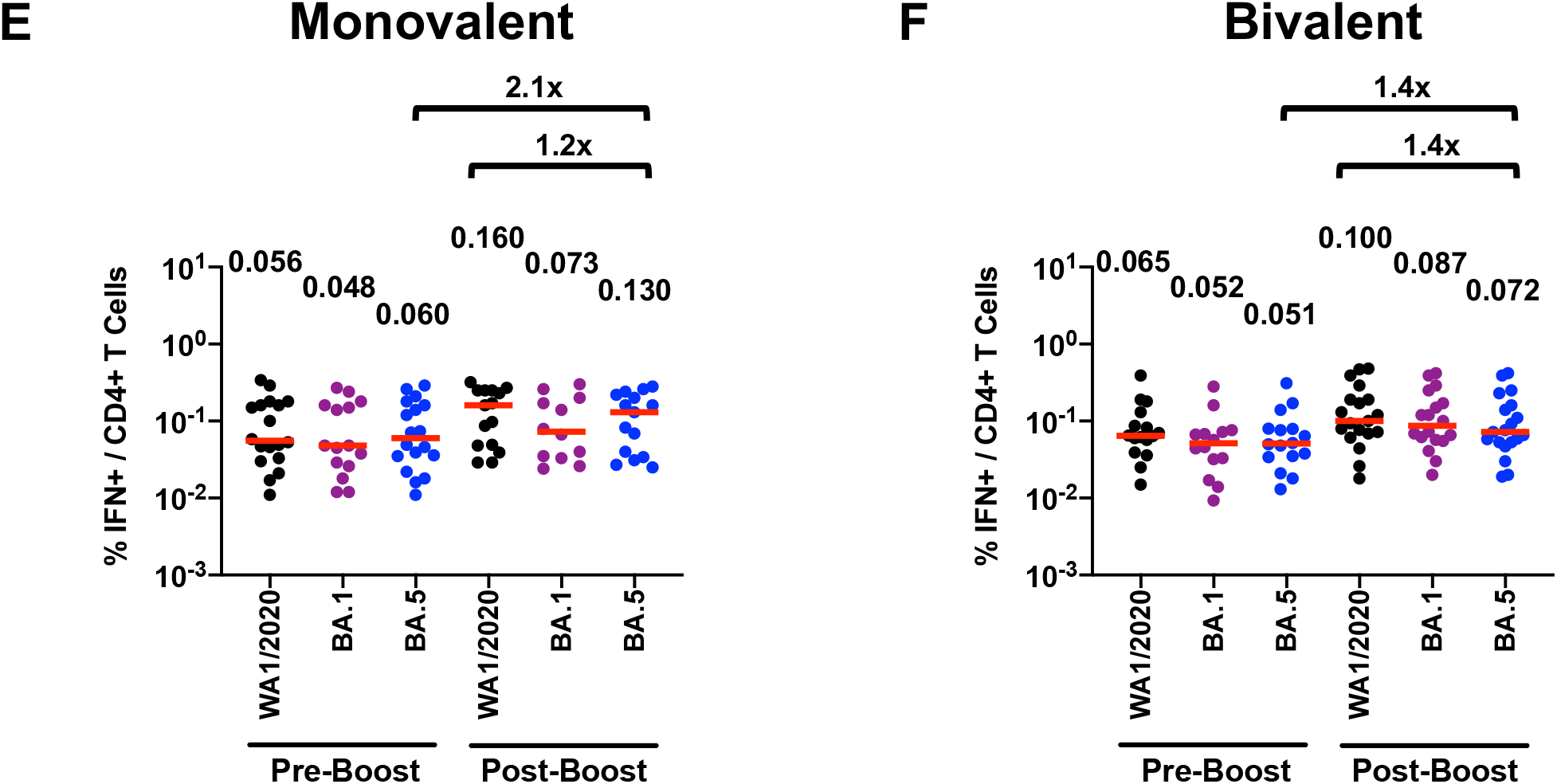
Neutralizing antibody and T cell responses following monovalent and bivalent mRNA boosting. **A, B.** Neutralizing antibody (NAb) titers by luciferase-based pseudovirus neutralization assays in individuals following monovalent or bivalent mRNA boosting. **C, D**. Spike-specific CD8+ T cell responses by intracellular cytokine staining assays in individuals following monovalent or bivalent mRNA boosting. **E, F**. Spike-specific CD4+ T cell responses by intracellular cytokine staining assays in individuals following monovalent or bivalent mRNA boosting. Responses were measured against the SARS-CoV-2 WA1/2020, Omicron BA.1, BA.2, and BA.5 variants. Medians (red bars) are depicted and shown numerically with fold differences.

Spike-specific CD8+ and CD4+ T cell responses increased only modestly following monovalent and bivalent mRNA boosting. Median BA.5 CD8+ T cell responses increased from 0.027% to 0.048% following monovalent mRNA boosting and from 0.024% to 0.046% following bivalent mRNA boosting (**Fig. 1C, 1D**). Median BA.5 CD4+ T cell responses increased from 0.060% to 0.130% following monovalent mRNA boosting and from 0.051% to 0.072% following bivalent mRNA boosting (**Fig. 1E, 1F**). Median BA.5 memory B cell responses were 0.079% following monovalent mRNA boosting and 0.091% following bivalent mRNA boosting (**Fig. S4**).

Our data demonstrate that both monovalent and bivalent mRNA boosters markedly increased antibody responses but did not substantially augment T cell responses. BA.5 NAb titers were comparable following monovalent and bivalent mRNA boosters, with a modest and nonsignificant trend favoring the bivalent booster by a factor of 1.3. These findings are consistent with data recently reported for a BA.1-containing bivalent mRNA booster^4^. Our findings suggest that immune imprinting by prior antigenic exposure^5^ may pose a greater challenge than currently appreciated for inducing robust immunity to SARS-CoV-2 variants.

## Data sharing

A.Y.C. and D.H.B. had full access to all the data in the study and take responsibility for the integrity of the data and the accuracy of the data analysis. All data are available in the manuscript or the supplementary material. Correspondence and requests for materials should be addressed to D.H.B.

## Supplementary Appendix

## Supplementary Methods

### Study population

A specimen biorepository at Beth Israel Deaconess Medical Center (BIDMC) obtained samples from individuals who received SARS-CoV-2 vaccines as well as monovalent or bivalent mRNA boosters. The BIDMC institutional review board approved this study (2020P000361). All participants provided informed consent. This study included 15 individuals who received the original monovalent mRNA boosters between January and August 2022 and 18 individuals who received the bivalent mRNA boosters in September 2022. Participants were excluded from this analysis if they were using immunosuppressive medications.

### Pseudovirus neutralizing antibody assay

Neutralizing antibody (NAb) titers against SARS-CoV-2 variants utilized pseudoviruses expressing a luciferase reporter gene^1^. In brief, the packaging construct psPAX2 (AIDS Resource and Reagent Program), luciferase reporter plasmid pLenti-CMV Puro-Luc (Addgene), and Spike protein expressing pcDNA3.1-SARS-CoV-2 SΔCT were co-transfected into HEK293T cells (ATCC CRL_3216) with lipofectamine 2000 (ThermoFisher Scientific). Pseudoviruses of SARS-CoV-2 variants were generated using the Spike protein from WA1/2020 (Wuhan/WIV04/2019, GISAID accession ID: EPI_ISL_402124), Omicron B.1.1.529 BA.1 (GISAID ID: EPI_ISL_7358094.2), BA.2 (GISAID ID: EPI_ISL_6795834.2), or BA.5 (GISAID ID: EPI_ISL_12268495.2). The supernatants containing the pseudotype viruses were collected 48h after transfection, and pseudotype viruses were purified by filtration with 0.45-μm filter. To determine NAb titers in human serum, HEK293T-hACE2 cells were seeded in 96-well tissue culture plates at a density of 2 × 10^4^ cells per well overnight. Three-fold serial dilutions of heat-inactivated serum samples were prepared and mixed with 50 μl of pseudovirus. The mixture was incubated at 37 °C for 1 h before adding to HEK293T-hACE2 cells. After 48 h, cells were lysed in Steady-Glo Luciferase Assay (Promega) according to the manufacturer’s instructions. SARS-CoV-2 neutralization titers were defined as the sample dilution at which a 50% reduction (NT50) in relative light units was observed relative to the average of the virus control wells.

### Enzyme-linked immunosorbent assay (ELISA)

SARS-CoV-2 spike receptor-binding domain (RBD)-specific binding antibodies in serum were assessed by ELISA. 96-well plates were coated with 0.5 μg/mL of SARS-CoV-2 WA1/2020, B.1.617.2 Delta, B.1.1.529 Omicron BA.1, BA.2, or BA.5 RBD protein in 1× Dulbecco phosphate-buffered saline (DPBS) and incubated at 4 °C overnight. After incubation, plates were washed once with wash buffer (0.05% Tween 20 in 1× DPBS) and blocked with 350 μL of casein block solution per well for 2 to 3 hours at room temperature. Following incubation, block solution was discarded and plates were blotted dry. Serial dilutions of heat-inactivated serum diluted in Casein block were added to wells, and plates were incubated for 1 hour at room temperature, prior to 3 more washes and a 1-hour incubation with a 1:4000 dilution of anti– human IgG horseradish peroxidase (HRP) (Invitrogen, ThermoFisher Scientific) at room temperature in the dark. Plates were washed 3 times, and 100 μL of SeraCare KPL TMB SureBlue Start solution was added to each well; plate development was halted by adding 100 μL of SeraCare KPL TMB Stop solution per well. The absorbance at 450 nm, with a reference at 650 nm, was recorded with a VersaMax microplate reader (Molecular Devices). For each sample, the ELISA end point titer was calculated using a 4-parameter logistic curve fit to calculate the reciprocal serum dilution that yields a corrected absorbance value (450 nm-650 nm) of 0.2. Interpolated end point titers were reported.

### Electrochemiluminescence assay (ECLA)

ECLA plates (MesoScale Discovery SARS-CoV-2 IgG Catalog No. K15463U-2 Panel 22 and Catalog No. K15575U-2 Panel 24) were designed and produced with up to 10 antigen spots in each well^2^. The plates were blocked with 50 uL of Blocker A (1% BSA in distilled water) solution for at least 30 minutes at room temperature shaking at 700 rpm with a digital microplate shaker. During blocking the serum was diluted to 1:5,000 in Diluent 100. The calibrator curve was prepared by diluting the calibrator mixture from MSD 1:9 in Diluent 100 and then preparing a 7-step 4-fold dilution series plus a blank containing only Diluent 100. The plates were then washed 3 times with 150 μL of Wash Buffer (0.5% Tween in 1x PBS), blotted dry, and 50 μL of the diluted samples and calibration curve were added in duplicate to the plates and set to shake at 700 rpm at room temperature for at least 2 h. The plates were again washed 3 times and 50 μL of SULFO-Tagged anti-Human IgG detection antibody diluted to 1x in Diluent 100 was added to each well and incubated shaking at 700 rpm at room temperature for at least 1 h. Plates were then washed 3 times and 150 μL of MSD GOLD Read Buffer B was added to each well and the plates were read immediately after on a MESO QuickPlex SQ 120 machine. MSD titers for each sample was reported as Relative Light Units (RLU) which were calculated using the calibrator.

### Intracellular cytokine staining (ICS) assay

CD4+ and CD8+ T cell responses were quantitated by pooled peptide-stimulated intracellular cytokine staining (ICS) assays^3^. Peptide pools contained 15 amino acid peptides overlapping by 11 amino acids spanning the SARS-CoV-2 WA1/2020, Omicron BA.1, or BA.5 Spike proteins (21st Century Biochemicals). 10^6^ peripheral blood mononuclear cells were re-suspended in 100 μL of R10 media supplemented with CD49d monoclonal antibody (1 μg/mL) and CD28 monoclonal antibody (1 μg/mL). Each sample was assessed with mock (100 μL of R10 plus 0.5% DMSO; background control), peptides (2 μg/mL), and/or 10 pg/mL phorbol myristate acetate (PMA) and 1 μg/mL ionomycin (Sigma-Aldrich) (100μL; positive control) and incubated at 37°C for 1 h. After incubation, 0.25 μL of GolgiStop and 0.25 μL of GolgiPlug in 50 μL of R10 was added to each well and incubated at 37°C for 8 h and then held at 4°C overnight. The next day, the cells were washed twice with DPBS, stained with aqua live/dead dye for 10 mins and then stained with predetermined titers of monoclonal antibodies against CD279 (clone EH12.1, BB700), CD4 (clone L200, BV711), CD27 (clone M-T271, BUV563), CD8 (clone SK1, BUV805), CD45RA (clone 5H9, APC H7) for 30 min. Cells were then washed twice with 2% FBS/DPBS buffer and incubated for 15 min with 200 μL of BD CytoFix/CytoPerm Fixation/Permeabilization solution. Cells were washed twice with 1X Perm Wash buffer (BD Perm/WashTM Buffer 10X in the CytoFix/CytoPerm Fixation/ Permeabilization kit diluted with MilliQ water and passed through 0.22μm filter) and stained with intracellularly with monoclonal antibodies against Ki67 (clone B56, BB515), IL21 (clone 3A3-N2.1, PE), CD69 (clone TP1.55.3, ECD), IL10 (clone JES3-9D7, PE CY7), IL13 (clone JES10-5A2, BV421), IL4 (clone MP4-25D2, BV605), TNF-α (clone Mab11, BV650), IL17 (clone N49-653, BV750), IFN-γ (clone B27; BUV395), IL2 (clone MQ1-17H12, BUV737), IL6 (clone MQ2-13A5, APC), and CD3 (clone SP34.2, Alexa 700) for 30 min. Cells were washed twice with 1X Perm Wash buffer and fixed with 250μL of freshly prepared 1.5% formaldehyde. Fixed cells were transferred to 96-well round bottom plate and analyzed by BD FACSymphony™ system. Data were analyzed using FlowJo v9.9.

### RBD-specific B cell staining

PBMCs were stained^4^ with Aqua live/dead dye for 20 min, washed with 2% FBS/DPBS buffer, and cells were suspended in 2% FBS/DPBS buffer with Fc Block (BD) for 10 min, followed by staining with monoclonal antibodies against CD45 (clone HI30, BV786), CD3 (clone UCHT1, APC-R700), CD16 (clone 3G8, BUV496), CD14 (clone M5E2, BV605), CD19 (clone SJ25C, BUV615), CD20 (clone 2H7, PE-Cy5), IgM (clone G20-127, BUV395), IgD (clone IA6-2, PE), IgG (clone G18-145, BUV737), CD95 (clone DX2, BV711), CD27 (clone M-T271, BUV563), CD21 (clone B-ly4, PE-CF594), CD38 (clone HB7, BUV805), and CD71 (clone M-A712, BV750). Staining with SARS-CoV-2 antigens including biotinylated WA1/2020 RBD (Sino Biological) labeled with FITC or BV605 or biotinylated BA.5 RBD (Sino Biological) labeled with APC or Alexa Flour605, and double positive B cells were considered RBD-specific. Staining was done at 4 °C for 30 min. After staining, cells were washed twice with 2% FBS/DPBS buffer, followed by incubation with BV650 streptavidin (BD Pharmingen) for 10 min, then washed twice with 2% FBS/DPBS buffer. After staining, cells were washed and fixed by 2% paraformaldehyde. All data were acquired on a BD FACSymphony flow cytometer. Subsequent analyses were performed using FlowJo software (BD Bioscience, v10.8.1). For analyses, in singlet gate, dead cells were excluded by Aqua dye and CD45 was used as a positive inclusion gate for all leukocytes. Within class-switched B cell population gated as CD19+ IgG+ IgM- IgD- CD3- CD14- CD16-, SARS-CoV-2 (WA1/2020) RBD-specific B cells were identified. The SARS-CoV-2-specific B cells were further distinguished according to CD21/CD27 phenotype distribution.

**Table S1.** Study population

|  | **Bivalent mRNA**<br>**booster**<br>**N=18** | **Monovalent mRNA**<br>**booster**<br>**N=15** |
| --- | --- | --- |
| **Age** (years), median (range) | 42 (37-57) | 50 (33-64) |
| **Sex at birth**, Female | 11 (61) | 13 (87) |
| **Race** |  |  |
| White | 16 (89) | 10 (67) |
| Asian | 2 (11) | 2 (13) |
| Black | 0 | 2 (13) |
| More than one race | 0 | 0 |
| Other* | 0 | 1 (7) |
| **Ethnicity** |  |  |
| Hispanic or Latino | 0 | 1 (7) |
| Non-Hispanic | 17 (94) | 14 (93) |
| Unknown | 1 (6) | 0 |
| **Medical condition** |  |  |
| Hypertension | 2 (11) | 3 (20) |
| Diabetes | 1 (6) | 1 (7) |
| Pregnant | 1 (6) | 2 (13) |
| Asthma | 2 (11) | 0 |
| **Most recent COVID-19 vaccine booster** |  |  |
| Pfizer monovalent booster | N/A | 5 (33) |
| Moderna monovalent booster | N/A | 10 (67) |
| Pfizer bivalent booster | 8 (44) | N/A |
| Moderna bivalent booster | 10 (53) | N/A |
| **Prior COVID-19 vaccine history** |  |  |
| BNT (4 doses) | 2 (11) | 0 |
| BNT (3 doses) | 8 (44) | 2 (13) |
| BNT / BNT / 1273 (3 doses) | 2 (11) | 1 (7) |
| BNT / BNT / Ad26 (3 doses) | 1 (6) | 4 (27) |
| 1273 (3 doses) | 3 (17) | 6 (40) |
| 1273 / BNT (2 doses) | 0 | 1 (7) |
| Ad26 / BNT / 1273 (3 doses) | 1 (6) | 0 |
| Ad26 / 1273 (2 doses) | 1 (6) | 1 (7) |
| **Days from last vaccine dose to sampling** | 21 (16-23) | 32 (17-64) |
| **Known COVID-19 positive test** | 6 (33) | 5 (33) |
| **Days from positive test to last vaccine\*\*** | 166 (149-257) | 143 (103-183) |

BNT=BNT162b2 (Pfizer); 1273=mRNA-1273 (Moderna); Ad26=Ad26.COV2.S (Janssen)
Data displayed as median (range or interquartile range, IQR) and n (%); BMI, body mass index; PCR, polymerase chain reaction; pregnant designation reflects time of last vaccine dose and/or time of sampling. All individuals with known prior infection had mild disease.
\*Self-reported race as Latina
\*\*Reported for only those with known prior infection

## Supplementary Figure Legends

**Figure S1.**
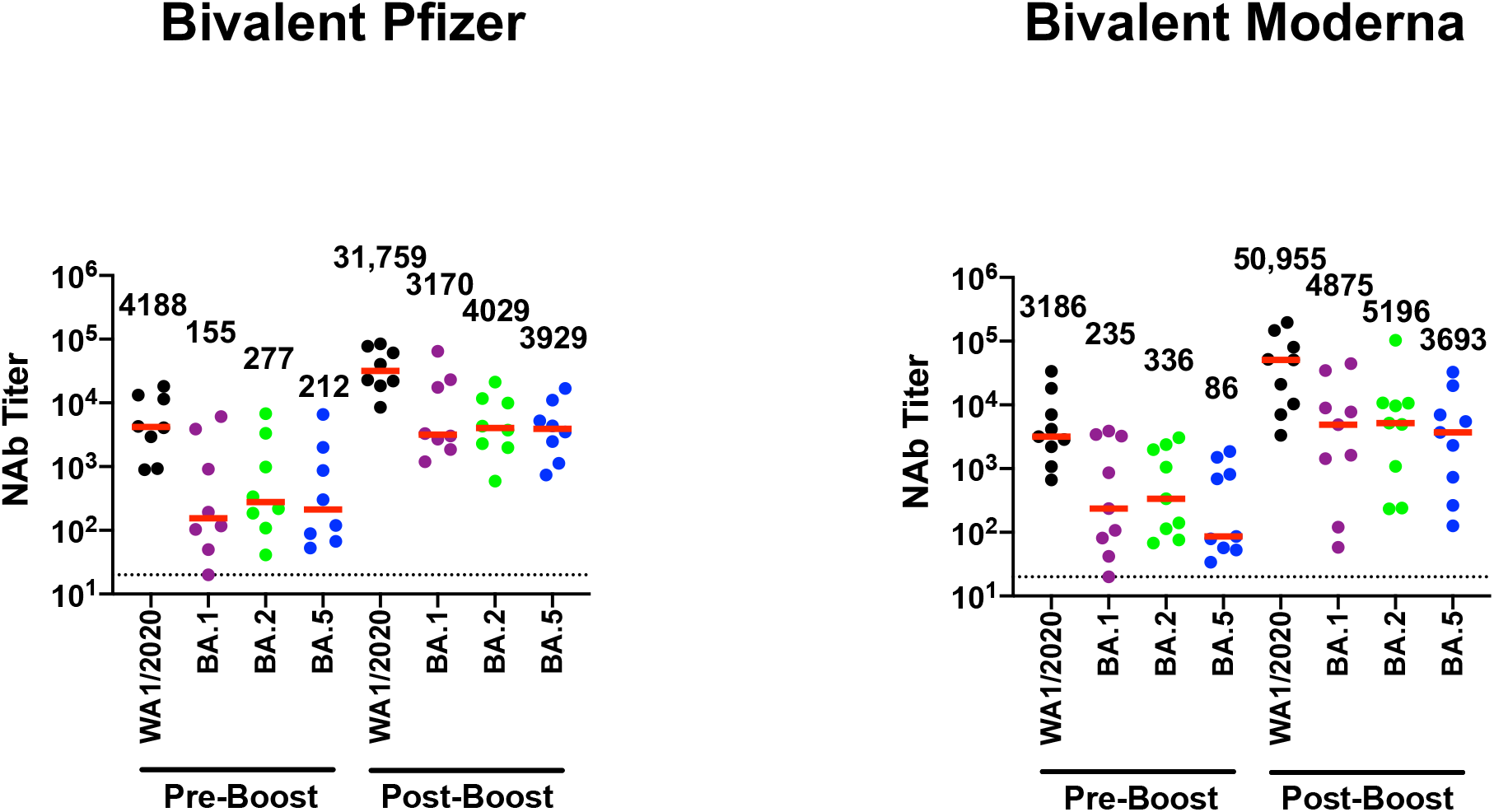
Neutralizing antibody responses following Pfizer or Moderna bivalent mRNA boosting. Neutralizing antibody (NAb) titers by luciferase-based pseudovirus neutralization assays in individuals following Pfizer or Moderna bivalent mRNA boosting. Responses were measured against the SARS-CoV-2 WA1/2020, Omicron BA.1, BA.2, and BA.5 variants. Medians (red bars) are depicted and shown numerically with fold differences.

**Figure S2.**
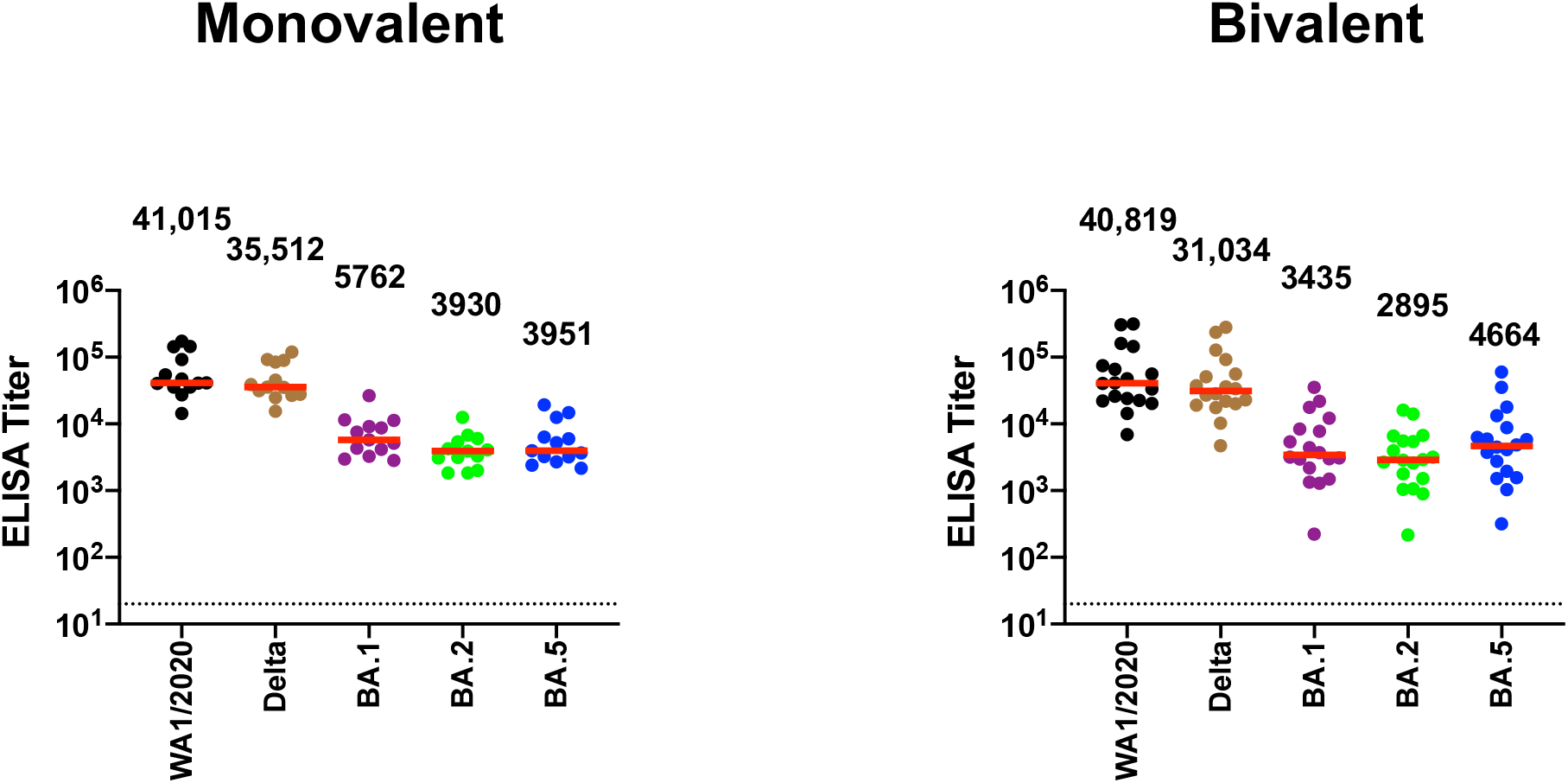
Binding antibody responses following monovalent and bivalent mRNA boosting by ELISA. Binding antibody titers by ELISA assays in individuals following monovalent or bivalent mRNA boosting. Responses were measured against the SARS-CoV-2 WA1/2020, Delta, Omicron BA.1, BA.2, and BA.5 variants. Medians (red bars) are depicted and shown numerically with fold differences.

**Figure S3.**
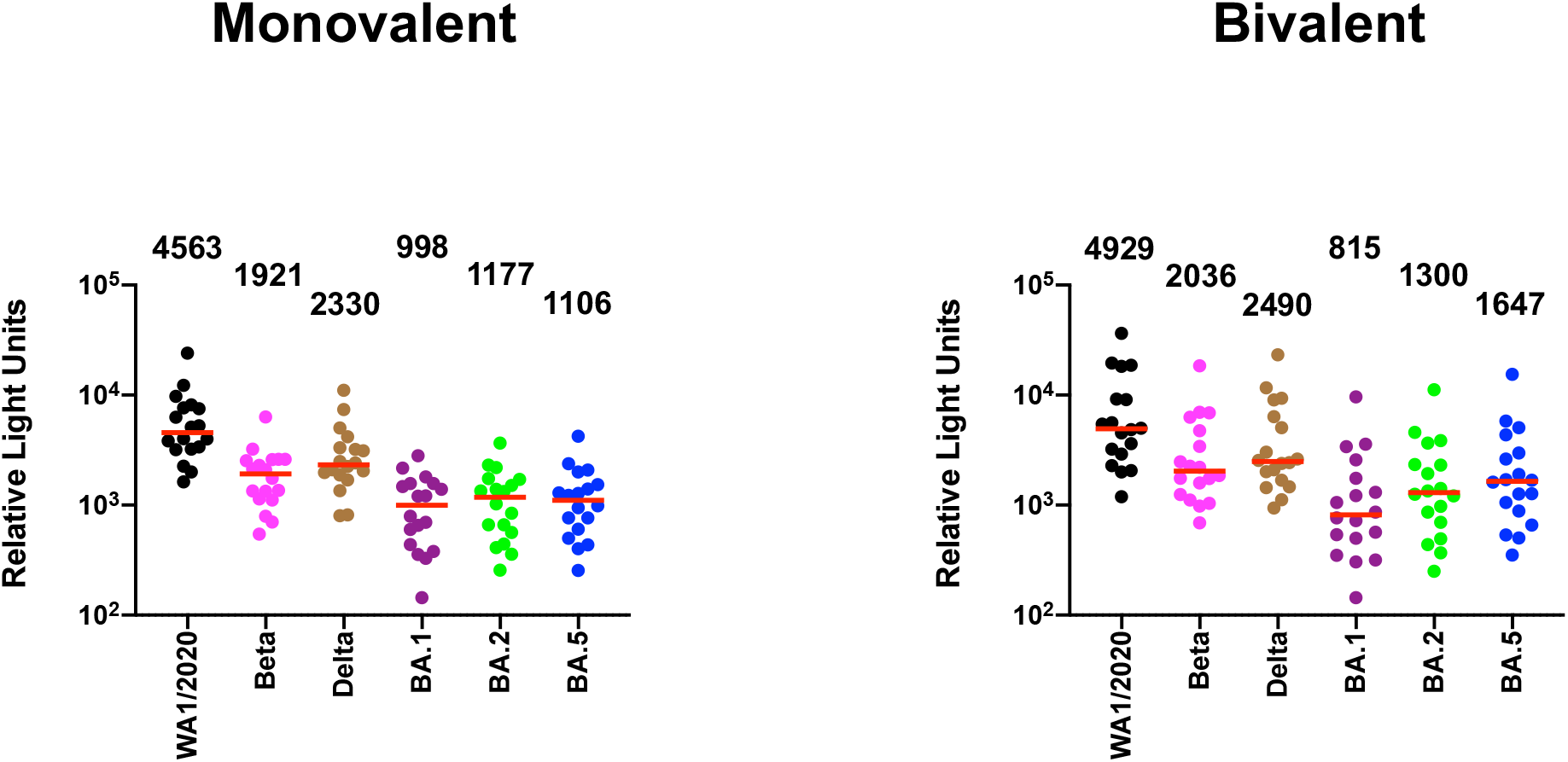
Binding antibody responses following monovalent and bivalent mRNA boosting by ECLA. Binding antibody titers by electrochemiluminescence (ECLA) assays in individuals following monovalent or bivalent mRNA boosting. Responses were measured against the SARS-CoV-2 WA1/2020, Beta, Delta, Omicron BA.1, BA.2, and BA.5 variants. Medians (red bars) are depicted and shown numerically with fold differences.

**Figure S4.**
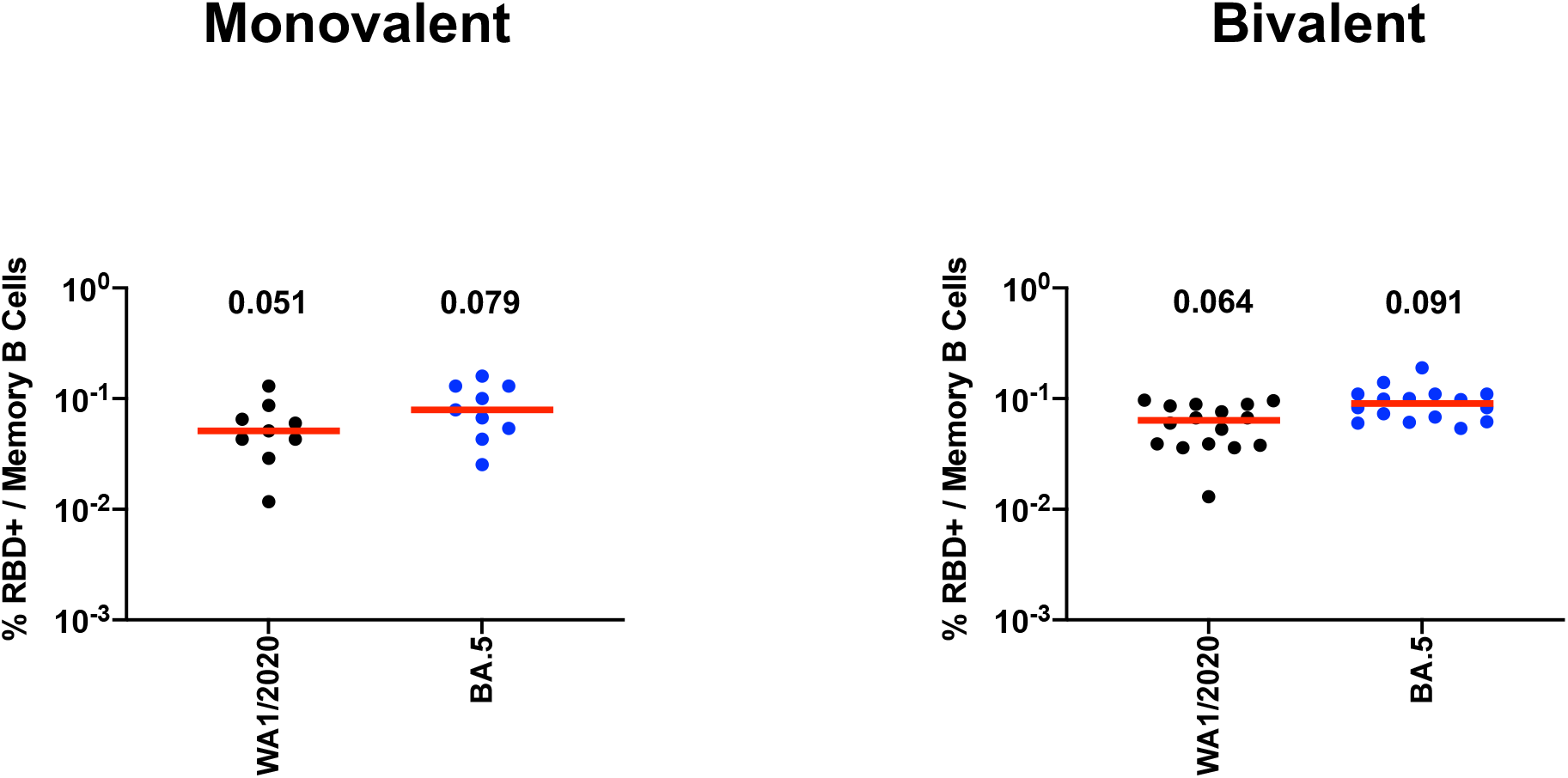
B cell responses following monovalent and bivalent mRNA boosting by flow cytometry. RBD-specific memory B cells by flow cytometry in individuals following monovalent or bivalent mRNA boosting. Responses were measured against the SARS-CoV-2 WA1/2020 and Omicron BA.5 variants. Medians (red bars) are depicted and shown numerically with fold differences.

